# Genomic loss of GPR108 disrupts AAV transduction in birds

**DOI:** 10.1101/2024.05.16.589954

**Authors:** Alexander A. Nevue, Anusha Sairavi, Samuel J. Huang, Hiroyuki Nakai, Claudio V. Mello

## Abstract

The G protein-coupled receptor 108 (*GPR108*) gene encodes a protein factor identified as critical for adeno-associated virus (AAV) entry into mammalian cells, but whether it is universally involved in AAV transduction is unknown. Remarkably, we have discovered that *GPR108* is absent in the genomes of birds and in most other sauropsids, providing a likely explanation for the overall lower AAV transduction efficacy of common AAV serotypes in birds compared to mammals. Importantly, transgenic expression of human *GPR108* and manipulation of related glycan binding sites in the viral capsid significantly boost AAV transduction in zebra finch cells. These findings contribute to a more in depth understanding of the mechanisms and evolution of AAV transduction, with potential implications for the design of efficient tools for gene manipulation in experimental animal models, and a range of gene therapy applications in humans.

## Introduction

Adeno-associated virus (AAV) is a non-pathogenic virus that is used as a vector for gene manipulation in a variety of tissues and organisms, as well as for a variety of gene therapy applications ^1^. Identifying and characterizing host cell factors necessary for efficient AAV transduction and understanding their variations in vertebrate species may provide novel insights into the mechanisms and evolution of AAV transduction. This knowledge may then be leveraged for more efficient gene manipulation applications, as well as in the rational design of targeted therapeutics to challenging or rare cell types. The adeno-associated virus receptor (*AAVR*, also known as *KIAA0319L*) has been identified by a genome-wide perturbation screen as coding for a near-universal receptor for AAV entry ^2^, but *GPR108* and *TM9SF2* are also critical for AAV transduction in mammals ^3, 4^. However, the transduction efficiencies of AAV serotypes vary widely across cell types, tissues, and species, and the exact roles of specific AAV entry factors and their interactions with capsid variants are unclear. In songbirds like zebra finches, which are a choice model organism for studying vocal production and learning, brain plasticity, sex steroid actions and adult neurogenesis ^5^, previous studies used AAV vectors to manipulate neuronal activation ^6-8^ or gene expression ^9-11^, or to identify specific cell types ^12, 13^ within vocal control areas. Nonetheless, AAV brain transduction efficiency in this and other bird species is reportedly varied and generally far lower than in laboratory animals such as mice ^12-18^. To better understand how evolutionary changes in AAV entry factors may have affected AAV transduction efficiency, we conducted genomic and functional analyses of these factors in the avian lineage, focusing on the zebra finch.

## Results

While AAVR is present in zebra finch, we found it has only moderate conservation (67.75% amino acid identity) with the human ortholog. Its polycystic kidney disease (PKD) domains, however, which have critical interactions with the AAV capsid ^19^, showed higher conservation between zebra finch and human (80-92% amino acid conservation, Fig. 1A). This conservation suggests that the key domains for AAV transduction are conserved in zebra finch AAVR, though divergence in AAVR sequence has been proposed as linked to AAV transduction inefficiency in birds ^18^. Intriguingly, even though *GPR108* was the second most highly enriched gene in the screen that identified AAVR ^2^ and has been confirmed as critical for AAV transduction in mammals ^3, 4^, it was absent in genome databases of all avian and most non-avian (crocodilians, snakes, and lizards) sauropsids where its genomic region is available in NCBI. This includes zebra finch, whose genome assembly is currently one of the most complete among vertebrates ^20^, as well as the more basal chicken, noting that the immediately syntenic genes (*RDH8*/*TRIP10*/*SH2D3A*) were also missing in birds (Fig. 1B), and *GPR108* was also absent from avian transcriptome databases available in NCBI. In contrast, *GPR108* was present in the turtle lineage, with the same number of exons and similar predicted protein size as in humans. This pattern suggests independent losses in different sauropsid lineages rather than a single loss in a common ancestor to all sauropsids, the ancestral loss in archosaurs (alligators and birds) followed by additional loss(es) of flanking genes in birds. All other genes enriched in the AAV entry factor screen were present in the zebra finch genome, and had similar predicted peptide lengths and high sequence conservation with their mammalian orthologs (Fig. 1C). The top genes from AAV transduction screens in mammalian cells, including *AAVR* and *TM9SF2*, showed broad expression throughout the zebra finch brain, including the striatum as well as auditory and vocal areas within the nidopallium and arcopallium (Fig. 1D), which are broadly considered analogous to superficial and deep layers respectively of mammalian auditory and motor cortices ^21-23^. These regions contain specific nuclei involved in the perceptual processing, learning and vocal-motor encoding of learned vocal patterns, and collectively form an interconnected system whose study has yielded numerous insights into the neurobiology of vocal communication ^5^. Their high expression in a broad range of regions suggests that gene expression regulation of most known entry factors is not limiting AAV brain transduction in songbirds.

**Figure 1:**
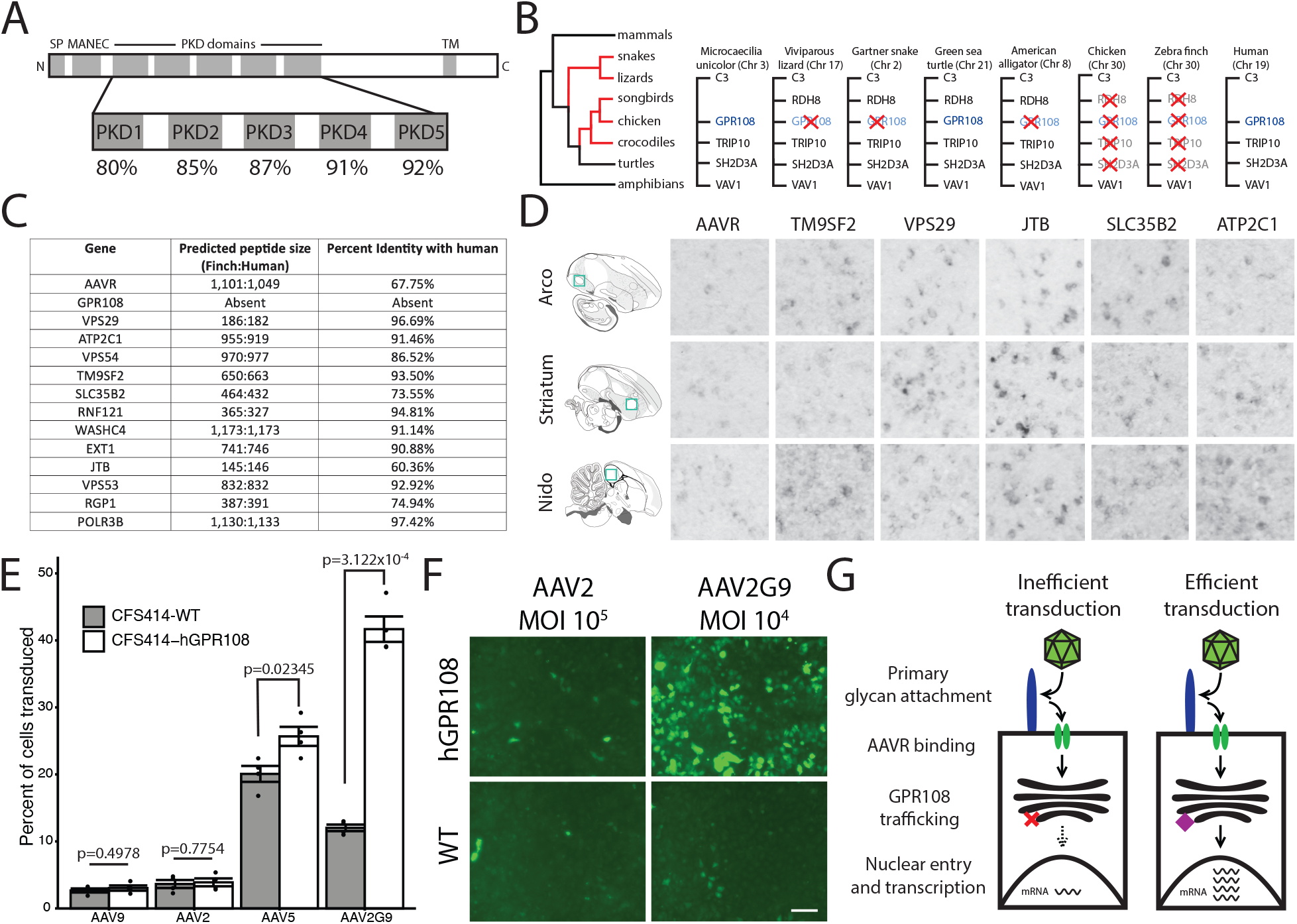
Genomic loss of GPR108 and link to AAV transduction effciency in songbirds. (A) Conservation of KIAA0319L (AAVR) PKD domains between human and zebra -finch. (B) Simpli-fied cladogram (left) depicting which amniote lineages lack GPR108 (in red), and diagrams showing the predicted GPR108 syntenic context of each lineage examined (right). (C) Comparison of the predicted peptide length and amino acid conservation between human and zebra -finch for the enriched genes in the AAV entry screen in Pillay et al., 2016 (D) Images of in situ hybridization of AAV entry factors in the zebra -finch brain. Green squares on sagittal sections on the left depict the areas shown on photomicrographs. The arcopallial (Arco) and nidopallium (Nido) areas shown are respectively analogous to deep motor cortical layers and super-ficial auditory cortical layers of mammals. These areas contain speci-fic auditory or vocal-motor nuclei of the zebra -finch song control circuitry. Images are 200 x 200 μm. (E) Quanti-fication of AAV transduction by FACS analysis in CFS414-WT and CFS414-hGPR108 cells. Shown is change in percent of cells transduced in GPR108+ compared to WT cells. P-values for comparisons from two-tailed unpaired t tests: AAV9: p=0.4978; AAV2: p=0.7754; AAV5: 0.02345; AAV2G9: 3.122x10-4. Data presented as means +/-SD, n=4 replicates per condition. (F) Images of GFP-immunoenhanced AAV transduction in CSF414-WT and CSF414-hGPR108 cells for AAV2 at MOI of 105 and AAV2G9 at MOI of 104. Scale bar is 50 μm. (G) Schematic representing main stages in AAV transduction; both surface glycan binding and GPR108-dependent signaling appear to be critical in zebra -finches.

To determine if the absence of *GPR108* impacts the efficiency of AAV transduction, we used a zebra finch cell line (CSF414) derived from embryonic fibroblasts ^24^ and generated transgenic zebra finch cells that express human *GPR108* (hGPR108). We then compared wild type (WT) and hGPR108+ cells for their ability to be transduced by AAV2, AAV9, AAV5 and AAV2G9.

Whereas AAV2 and AAV9 and known to be GPR108-dependent, AAV5 is GPR108-independent and therefore more likely to transduce a cell type lacking GPR108 ^3^. AAV2G9 ^25^ is a chimeric capsid which possesses both galactose and heparin binding domains of AAV9 and AAV2. Both AAV2 and AAV9 demonstrated similar transduction efficiencies between WT and hGPR108+ cells (Fig. 1E). AAV2, which robustly transduces cell lines expressing heparin sulfate proteoglycan ^26^, was unable to transduce zebra finch cells at a multiplicity of infection (MOI) of 10^5^ (Fig. 1F). In sharp contrast, AAV2G9 transduced hGPR108+ cells much more robustly than WT cells, even at a lower MOI of 10^4^ (Fig. 1F), providing evidence that robust AAV transduction is dependent on both the presence of GPR108 and the capacity to bind to glycans on the surface of zebra finch cells (Fig. 1G). We also observed higher AAV5 transduction of WT cells than was achieved by AAV2 or AAV9 (Fig. 1E), consistent with AAV5’s ability to transduce independently of GPR108 ^3^. Interestingly, we detected a small but significant enhancement of AAV5 transduction in hGPR108+ compared to WT cells (Fig. 1E), although the effect was much less pronounced than for AAV2G9.

To gain further insight into how a lack of *GPR108* may affect brain cell infectivity by common AAV serotypes, we used a barcoded capsid strategy that allows for a rigorous quantitative assessment of brain tropism of AAV serotypes. Injections of an RNA/DNA-barcoded AAV capsid library^27^ containing AAV serotypes 1-11 into striatal and arcopallial regions of zebra finches revealed that AAV9 produces the most robust transduction in both areas based on AAV transcript levels (Fig. 2A). Interestingly, the AAV biodistribution, reflected in relative DNA barcode reads, did not correlate with transduction efficiency, reflected in relative RNA barcode reads, as AAV8 and AAV7 genomes were found in more abundance than AAV9 genomes in both brain regions (Fig. 2B). This finding suggests that, while AAV8 and AAV7 may be more infectious and deliver more vector genome DNA copies to zebra finch brain tissue than other serotypes, their transduction may be prevented at the level of transcription or translation by the presence or absence of other regulatory factors in the zebra finch brain. We also compared the transduction of AAV9 to two other commonly used viral vectors, VSV G pseudotyped lentivirus and adenovirus serotype 5 (Ad5). We observed low transduction efficiency with VSV G pseudotyped lentivirus, consistent with our previous description of a truncated VSV G protein receptor (*LDLR*) in songbirds ^28^. We also observed poor transduction of Ad5, which has been effective for transgenesis in quail ^29^ (Fig. 2C), suggesting that Ad5 receptor entry factors may also be divergent between mammals and songbirds.

**Figure 2:**
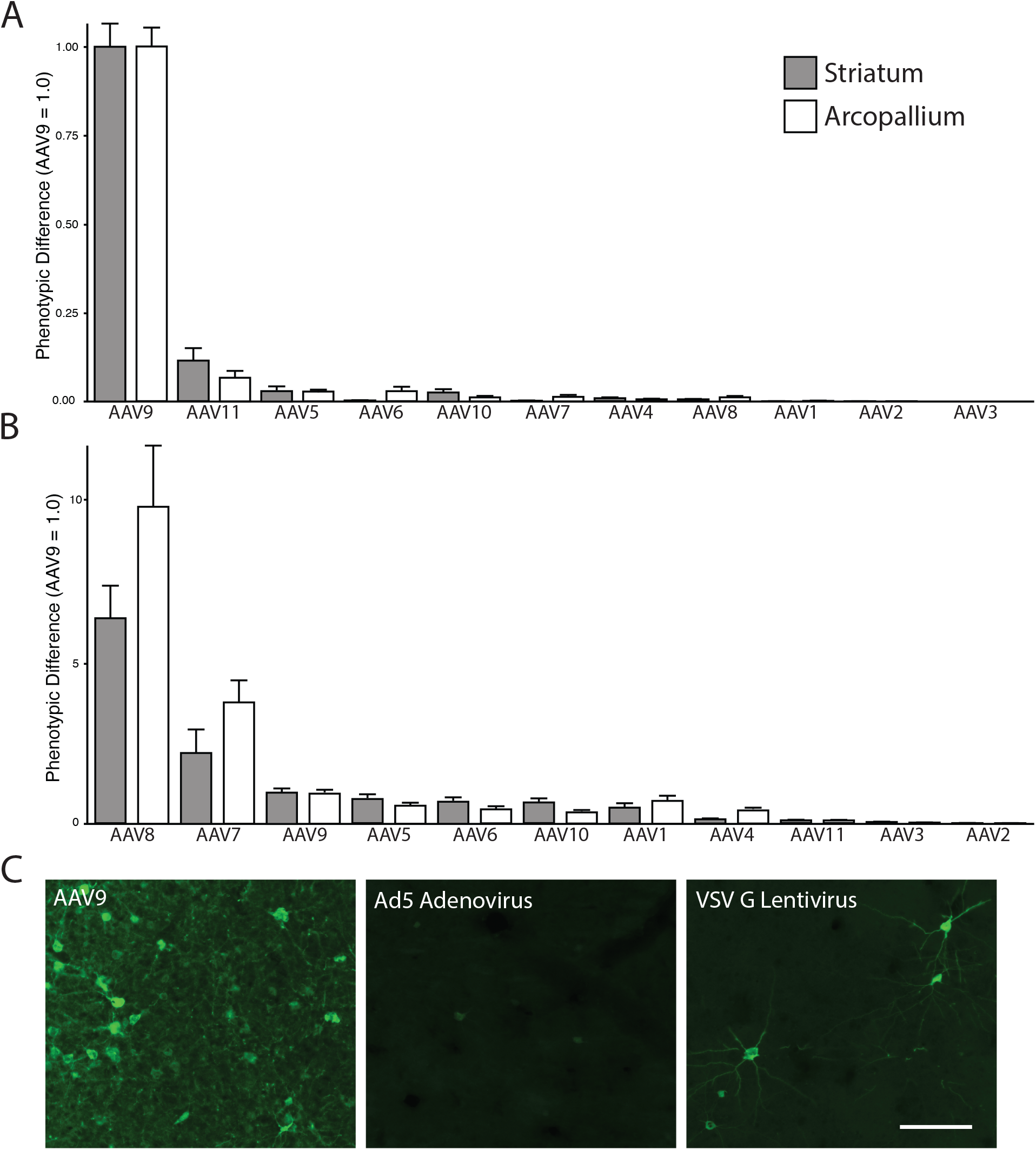
AAV Barcode-Seq reveals transduction effciencies of capsids in the zebra -finch brain. (A) Transduction effciency measured by AAV RNA Barcode-Seq. (B) Relative quantity of AAV vector genome DNA delivered by each capsid, measured by AAV DNA Barcode-Seq. For both B and C, phenotypic difference is normalized to AAV9. (C) Representative images of transduction effciency of AAV9, Ad5, and VSV G pseudotyped lentivirus vectors stereotaxically injected into the nidopallium of zebra -finches. Scale bar, 50 μm.

## Discussion

The absence of *GPR108* in the zebra finch genome can be interpreted as a naturally occurring gene knockout, and the ability of human *GPR108* to enhance the susceptibility of zebra finch cells to AAV transduction further supports a *GPR108* role in AAV transduction. The lack of *GPR108* extends to other bird groups, likely resulting from a genomic loss in an archosaur ancestor, as well as to other sauropsids. Strains of AAVs have been isolated from some of these species that lack *GPR108* ^30^, showing that *GPR108*-independent variants may have evolved, such as AAV5 ^3^. In the case of avian AAV (AAAV), it is highly divergent in sequence compared to AAV strains isolated from primates ^30^. Of note, AAAV uses galactose for cellular attachment, as we have also shown here, suggesting that surface glycan binding may be a critical requirement for transducing avian cells ^31^. Notably, there is incongruence between the transduction we observed *in vitro* and *in vivo*, as illustrated by the lack of AAV5 transduction observed *in vivo*.

This finding points to cell type- and tissue-specific transduction differences, extracellular glycans being a likely contributing factor. This interpretation is also consistent with enhanced AAV2G9 transduction in the zebra finch brain seen with modifications of the heparin binding domain ^13^. Together with our report on divergence of the avian homolog of the VSV G receptor *LDLR* ^28^, our present findings point to multiple losses of functional viral receptors in avian genomes, consistent with a complex genomic landscape for host-viral interactions of relevance for emerging zoonotic viruses ^32^.

While *AAVR* and *GPR108* have been identified as key AAV entry factors, they are not universal. AAV4 can enter cells independently of AAVR ^33^ and AAV5 can transduce cells independently of *GPR108* ^3^. Determining the mechanisms by which these capsids are able to transduce cells may offer alternative strategies for targeted therapeutics. Modifying the expression of entry factors is one possible mechanism for improving transduction efficiency, as recently shown by AAVR overexpression ^34^. However, as suggested by our barcoded capsid analysis, the relationship between the target cell type’s transcriptome and AAV capsid is also key, and can potentially be leveraged for improved transduction, a strategy that would serve as an alternative to directed evolution ^35^. In that context, our data highlight the importance of deep transcriptional and biochemical investigation of target cells for AAV mediated gene therapy. Rapidly developing single cell and spatial genomics technology ^36^ may further elucidate cell type- and capsid-specific entry factors. The brain is a particularly complex tissue that contains many highly distinct regions, each with a multitude of neuronal and non-neuronal cell types. With many therapeutics relying on efficient delivery to specific cell types, and the probable regional diversity in cell surface attachment factors and expression levels of entry factors, the development of engineered AAVs for effective brain gene therapy must take a wholistic perspective on both genomic and transcriptomic components.

## Methods

### Animals

All procedures involving live animals were approved by the Institutional Animal Care and Use Committee at Oregon Health & Science University and in accordance with NIH guidelines. Adult zebra finches (Taeniopygia guttata) were obtained from our colony or purchased from a local breeder.

### Genomics

For a thorough and systematic examination of *GPR108* in birds and other sauropsids, we started by searching databases (NCBI’s RefSeq, Ensembl) for genes annotated as *GPR108* or described as “G protein-coupled receptor 108”, which revealed *GPR108* in turtles only. Orthology of GPR108 in turtles to human *GPR108* was confirmed by reciprocal BLAST alignments and synteny verification. We then used turtle *GPR108* RefSeq mRNA sequences as queries for low similarity BLAST searches in sauropsid NCBI’s genomic (RefSeq_genome or _rna, whole genome shotgun) and transcriptome (Transcriptome Shotgun Assembly, EST) databases, which yielded no significant hits besides *GPR107*, a related gene. We next examined closely the syntenic region where GPR108 is expected to occur. We focused on four songbirds (*Taeniopygia guttata, Catharus ustulatus, Corvus moneduloides, Pseudopodoces humilis*), a falcon (*Falco rusticolus*), a pigeon (*Columba livia*), a duck (*Anas platyrhynchos*), and chicken (*Gallus gallus*), compared to representative alligator (*Alligator mississippiensis*), turtle (*Chelonia mydas*), snake (*Thamnophis elegans*), lizard (*Zootoca vivipara*), and amphibian (*Microcaecilia unicolor*) species, as well as mammals (*Homo sapiens, Mus musculus*). The avian species chosen were the only ones where the region containing *C3* and *VAV1* is well assembled, noting that these genes flank the region containing GPR108 and/or its immediately syntenic genes (*RDH8*/*TRIP10*/*SH2D3A*) in other vertebrates (non-avian sauropsids, mammals, amphibians). This region is within avian microchromosome 30, which has been very difficult to fully sequence and assemble in birds, typically requiring long-read PacBio technology ^20^. We verified that *C3* and *VAV1* are present and contiguous in all birds examined, with no intervening sequence gaps. For the *AAVR* analysis, the sequences of *KIAA0319L* from zebra finch (Gene ID: 100217975) and human (Gene ID: 79932) were imported into InterProScan for domain prediction, and MUSCLE was used for sequence alignments.

### In situ hybridization

*In situ* hybridizations for *AAVR* (NCBI cDNA clone CK304065), *TM9SF2* (FE735586), *VPS29* (CK304285), *JTB* (FE721652), *SLC35B2* (CK316607), and *ATP2C1* (DV948245) were carried out as previously described ^21, 37^. Briefly, plasmids containing cDNA of the gene of interest were isolated and restriction enzyme digested with BSSHII to release the insert. The insert was purified using a QIAquick PCR purification kit (Qiagen). Antisense DIG-tagged probes were synthesized by *in vitro* transcription using T3 polymerase (Promega) and purified using a Sephadex G-50 column. Probes were hybridized to brain sections from adult male zebra finches overnight at 65°C and washed at high stringency. Sections were blocked an incubated in anti-DIG-AP Fab fragments (1:600; Roche) for two hours and incubated in BCIP/NBT chromogen (PerkinElmer) overnight. Probe specificity was examined by aligning the corresponding sequences to the zebra finch genome and verifying that alignment was specific to the expected loci. As further specificity controls, the patterns seen in replicates (n=3 birds) were compared with negative controls where riboprobes were omitted, and with a broad range of genes with known expression patterns in the zebra finch brain ^38^.

### Cell Culture

The cell line CFS414 derived from zebra finch embryonic fibroblasts^24^ was obtained as a gift from the Jarvis lab (the Rockefeller University). Cells were cultured in DMEM supplemented with 10% FBS and 1% each of L-Glutamine and Penicillin-Streptomycin. To generate the transgenic line expression hGPR108, 1.9 µg of the transposon (pSBbi-RP, Addgene plasmid #60513) ^39^ and 100 ng of the transposase (pCMV(CAT)T7-SB100, Addgene plasmid #34879) ^40^ were transfected using polyethyleneimine (PEI) with a ratio DNA:PEI of 1:2. One day after transfection, the cells were subjected to selection with 1 µg/mL puromycin and selection continued for 5 days until all cells were RFP+. Human GPR108 (NCBI RefSeq NP_001073921) was synthesized as GeneArt Strings by Invitrogen and cloned into pSBbi using Gibson cloning.

### AAV particle production

Double-stranded AAV vectors carrying a CMV promoter-driven GFP expression cassette were produced in HEK293 cells using an adenovirus-free triple transfection method. Briefly, cells seeded in T225cm^2^ flasks were transfected with the DNA plasmids using PEI at a DNA:PEI ratio of 1:2. At 5 days post-transfection, cells and media were harvested and purified by two cycles of cesium chloride ultracentrifugation followed by 4 rounds of dialysis, as previously described ^41^.

### In vitro transduction and FACS analysis

CFS414-WT and CFS414-hGPR108 cells were seeded 24 hours prior to infection with AAV9, AAV2, AAV5, or AAV2G9, each expressing GFP, at an MOI of 10^5^ in the absence of a helper virus. This experiment was performed in quadruplicates. Four days after infection, the percent of GFP+ cells was determined using a BD FACSymphony Analyzer and FlowJo. Two-tailed unpaired *t* tests were performed using R. In a separate experiment, AAV2 at a MOI of 10^5^ and AAV2G9 at an MOI of 10^4^ were introduced to CFS414-WT and CFS414-hGPR108 cells in the absence of a helper virus. After two days, cells were fixed with 4% paraformaldehyde (PFA) and GFP+ transduction was immunoenhanced using an anti-GFP 488 antibody (ThermoFisher).

### AAV Barcode-Seq and histological analysis

The barcoded library pdsAAV-U6-VBCx ^41^ (250 nL) was stereotaxically injected into either the arcopallium (targeting song nucleus RA) or the striatum (targeting song nucleus Area X) of adult male zebra finches (n=2). After two weeks, finches were sacrificed via decapitation and the regions of interest were grossly dissected and flash frozen in a dry ice/isopropyl alcohol slurry.

Dissected tissue was mixed and split into two parts. One part was homogenized in Trizol using a bead mill homogenizer and RNA was extracted using chloroform phase separation. DNA was extracted from the second part using a Qiagen DNeasy Blood and Tissue kit. AAV Barcode-Seq samples were processed as previously described ^27^. Briefly, RNA was reverse transcribed using a High-Capacity cDNA Reverse Transcription Kit (Applied Biosciences). Viral barcodes (VBCs) were PCR amplified and indexed using sample-specific barcodes (SBCs). The amplicons were mixed at an equimolar ratio and sequenced on an Illumina NovaSeq 6000. Phenotypic difference was determined by quantifying the Illumina sequence read numbers for each VBC and normalizing to the library stock ^27^.

For single virus injections, 250 nL of each virus was injected into the arcopallium of a female zebra finch. AAV9-hSyn-eGFP was prepared as described above and had a titer of 7.1x10^12^ vg/mL. HDAd5-hSyn-eGFP adenovirus was acquired from the University of Iowa Viral Vector Core with a titer of 6.3x10^10^ IGU/mL. VSV G pseudotyped lentivirus was prepared at the OHSU virology core using pHR-hSyn-eGFP plasmid (Addgene plasmid #114215) with a titer of 1.0x10^9^ TU/mL. After two weeks, finches were perfused first with 0.9% NaCl followed by 3% PFA. Brains were dissected and post-fixed in 3% PFA overnight and transferred to 30% sucrose for 2 days.

Brains were then frozen in Tissue-Tek (Sakura) and sectioned at 30 µm on a cryostat. Immunohistochemistry was performed to enhance the GFP signal. Overnight, free floating sections were incubated in TNB (100 mM Tris-HCl pH 7.4, 150 mM NaCl, 0.36% w/v BSA, 1% skim milk), 0.3% Triton X-100 (Sigma), and anti-GFP 1:1000 (Abcam, ab13970). The next day, sections were washed and incubated in TNB, 0.3% Triton X-100, and Alexa Fluor 488-labeled goat anti-chicken IgG 1:1000 (Invitrogen, A11039) for two hours. Sections were then washed, mounted on ColorFrost Plus slides (Fisher), and imaged using a Keyence BZ-X710 fluorescence microscope.

## Data availability

Raw data and CFS414-hGPR108 cell line are available upon request to the corresponding authors.

## Acknowledgments

This work was supported by NIH R01NS088399 to HN, R21OD028774 to CVM, and F32MH134413 to AAN. We thank Matthew Biegler and Erich Jarvis for providing the CFS414 cell line. We also thank Samuel Young for advice on the use of Ad5.

## Author contributions

AAN, AS, SJH, HN, and CVM conceived of the study. AAN and CVM performed the genomics analyses. AAN and SJH performed the Barcode-Seq analysis. AAN and AS performed the *in vitro* transduction studies. AAN analyzed the data. AAN wrote the manuscript with input from HN and CVM. All authors edited the manuscript.

## Competing interests

HN receives royalties of AAV-related technologies licensed by Takara Bio Inc. and Capsigen Inc., serves as a consultant for biotech companies, and is a co-founder of Capsigen Inc., a company commercializing AAV vector-related technologies. SJH receives royalties from Capsigen Inc. All other authors declare no competing interests.

